# A sheath of motile cells supports collective migration in of the Zebrafish posterior lateral line primordium under the skin

**DOI:** 10.1101/783043

**Authors:** Damian Dalle Nogare, Naveen Natesh, Ajay Chitnis

## Abstract

During embryonic development, cells must navigate through diverse three-dimensional environments robustly and reproducibly. The zebrafish posterior lateral line primordium (PLLp), a group of approximately 120 cells which migrates from the otic vesicle to the tip of the tail, spearheading the development of the lateral line sensory system, is an excellent model to study such collective migration in an in vivo context. This system migrates in a channel formed by the underlying horizontal myoseptum and somites, and the overlying skin. While cells in the leading part of the PLLp are flat and have a more mesenchymal morphology, cells in the trailing part progressively reorganize to form epithelial rosettes, called protoneuromasts. These epithelial cells extend basal cryptic lamellipodia in the direction of migration in response to both chemokine and FGF signals. In this study, we show that, in addition to these cryptic lamellipodia, the core epithelial cells are in fact surrounded by a population of motile cells which extend actin-rich migratory processes apposed to the overlying skin. These thin cells wrap around the protoneuromasts, forming a continuous sheath of cells around the apical and lateral surface of the PLLp. The processes extended by these cells are highly polarized in the direction of migration and this directionality, like that of the basal lamellipodia, is dependent on FGF signaling. Consistent with interactions of sheath cells with the overlying skin contributing to migration, removal of the skin stalls migration. However, this is accompanied by some surprising changes. There is a profound change in the morphology of the sheath cells, with directional superficial lamellipodia being replaced with the appearance of undirected blebs or ruffles. Furthermore, removal of the skin not only affects underlying lamellipodia, it simultaneously alters the morphology and behavior of the deeper basal cryptic lamellipodia, even though these cells do not directly contact the skin. Directional actin-rich protrusions on both the apical and basal surface and migration are completely and simultaneously restored upon regrowth of the skin over the PLLp. We suggest that this system utilizes a circumferential sheath of motile cells to allow the internal epithelial cells to migrate collectively in the confined space of the horizontal myopseptum and that elastic confinement provided by the overlying skin is essential for effective collective migratory behavior of primordium cells.

## Introduction

Collective cell migration is fundamental for embryonic development^1^ and its dysregulation during morphogenesis is a key contributor to many developmental disorders^2^. Recent studies have additionally placed collective cell migration at the heart of metastasis of certain types of cancer^3,4^. These physiological and pathogenic contexts are linked by the necessity for cells to migrate, often through diverse 3D environments, while maintaining group cohesion and directionality.

In recent years, the Zebrafish posterior lateral line primordium (PLLp) has emerged as a powerful model for studying a wide range of cellular and developmental processes, including cell-cell signaling, tissue patterning, and collective migration^2,5,6^. This group of 100-150 cells is initially specified adjacent to the otic vesicle and migrates caudally down the length of the embryo over the course of ~24 hours. As this primordium migrates, cells in the tailing domain are progressively reorganized into apically-constricted epithelial rosettes, each cradling a central sensory hair cell progenitor^7^. Cells in the trailing-most rosette in the PLLp eventually lose the ability to sustain collective migration and are deposited by the migrating primordium. These deposited clusters will go on to develop into the mature sense organs of the lateral line, a sensory system that detects water flow about the animal^8^. Cells in the PLLp that are not incorporated into these epithelial rosettes are deposited as a continuous stream of so-called “interneuromast cells”, which lie between the sense organs^9,10^.

Migration of the PLLp is guided by a unidirectional stripe of a chemokine, Cxcl12a, secreted by muscle pioneer cells lying along the horizontal myoseptum^11,12^. Directional interpretation of this uniform stripe is accomplished by the expression of two chemokine receptors in the PLLp. The leading two-thirds of the PLLp express the chemokine receptor Cxcr4b, which can bind Cxcl12a and activate productive G-protein coupled signaling to drive cell migration. The trailing-most cells, on the other hand, express the Cxcr7b chemokine receptor, which can bind Cxcl12a but cannot activate G-protein coupled signaling and is thought to act as a sink for ligand^12–14^. Together, these two receptors generate a tissue-scale gradient of Cxcl12a availability along the length of the primordium which drives directional migration^15,16^, and provides an inherent polarity in the competence of cells to transduce the chemokine signal.

In cell transplantation experiments, basal cryptic lamellipodia are observed extending from PLLp cells in the direction of migration^14,17^, a common strategy for migrating epithelial cells^18^. Crucially, these lamellipodia are observed extending from both leading mesenchymal cells and trailing epithelial cells^14^, suggesting that cells along the length of the PLLp contribute to migration. This is consistent with recent studies showing that chemokine signaling is necessary along the entire *cxcr4b*-positive domain to support effective collective migration^19^. In addition to chemokine signaling, fibroblast growth factor (Fgf) signaling is also required for migration. The polarization of these basal migratory protrusions appears to be dependent on active Fgf signaling; their polarity is lost upon Fgf receptor inihbition, even when chemokine signaling is unperturbed, and this occurs concomitantly with a loss of migratory ability^17^. Furthermore, experiments with isolated PLLp fragments generated by laser ablation suggest that Fgf could act as a direct migratory cue^20^. These two systems, and potentially others, act together to govern collective migration of the PLLp.

Less studied, however, is the way in which the PLLp interacts with surrounding tissue as it migrates and what influence surrounding tissue might have on migration and morphogenesis. Aman *et al* showed that traversing underlying intersomitic boundaries does not influence the deposition of neuromasts, as the lateral line primordium does not deposit more closely spaced neuromasts in *trilobyte* mutants, which have more densely packed somites^21^. Other studies have shown that the directionality of migration does not rely on any extrinsic cues from the surrounding tissue and that directional migration is an autonomous property of the primordium itself^14^. However, the primordium has a dramatic effect on the tissue through which it migrates. The PLLp migrates along the horizontal myoseptum, between between the underlying somites and overlying skin. As it migrates, the skin is displaced upwards and is separated from the underlying tissue by the passage of the PLLp, returning rapidly to its original apposition with the underlying somites after the passage of the PLLp.

In this study, we focus on the PLLp cells that lie on the apical side of the PLLp, above the apical constrictions of the neuromasts, and make extensive contacts with the overlying skin. We show that these cells extend directional migratory processes and that the directionality of these processes, like that of the basal cryptic lamellipodia is dependent on Fgf signaling. Furthermore, we show that mechanically removing the skin prevents PLLp migration, which is subsequently recovered when the skin heals back over the PLLp. Loss of the overlying skin disrupts polarization of both apical and basal migratory processes. Taken together, these data suggest that the PLLp coordinates collective migration by extending lamellipodia both basally, against the underlying tissue, and apically, against the skin, and that confinement between the skin and underlying somites is essential for PLLp migration.

## Materials and Methods

### Fish Lines and Embryo Manipulation

Zebrafish embryos were generated by natural spawning, maintained under standard conditions, and staged according to Kimmel et al^22^. The following lines were used: *Tg(cldnb:lyn-gfp)*^14^, *TgBAC(cxcr4b:lifeact-citrine)*^23^, *TgBAC(cxcr4b:h2a-mcherry)*^19^, *Tg(cldnb:lyn-mscarlet)*, *Tg(krt4:dsred)*^24^. For FGF receptor inhibition experiments, embryos were treated with 20μM SU5402 (Tocris) 6 hours prior to imaging. For skin removal experiments, embryos were embedded in 2% low-melt agarose in Fluorobrite media. Agarose above the PLLp was removed with forceps, and a tungsten needle (Roboz surgical instruments) was used to manually remove the skin above the PLLp. Following skin removal, embryos were dissected from the agarose and re-embedded in 1% low-melt agarose in Fluorobrite for imaging.

For generation of *Tg(cldnb:lyn-mscarlet)* transgenic fish, the lyn-mscarlet^25^ DNA sequence was codon optimized for zebrafish expression^26^ and commercially synthesized. This fragment was cloned downstream of the 4.2kb *claudinb* promoter fragment^27^, which drives expression in the lateral line primordium and periderm, among other tissues. This construct was cloned between sites for the Tol1 transposon^28^ and 20ng of plasmid DNA was injected with 80ng of *tol1* mRNA into 1-cell stage zebrafish embryos. Founders were screened by fluorescence for high expression in the lateral line primordium.

For generation of chimeric PLLp, we dechorionated embryos at ~2hpf, and placed them in embryo media with 100 U/mL penicillin and 0.1mg/mL streptomycin (Roche). When the embryos had reached high-sphere stage (~3.5-4hpf), they were placed in individual wells made in agarose by a custom-printed mold. An Eppendorf CellTram Vario connected to a glass capillary needle with the tip removed at approximately the diameter of an embryonic cell was used to gently aspirate cells from the host embryo and place them in the donor embryo. After transplantation, embryos were placed in individual chambers of a 48-well plate in embryo medium with Penicillin and Streptomycin and grown overnight at 28°C. Embryos were screened at 24hpf for expression of the donor transgene in the PLLp.

### Time-lapse microscopy, segmentation, and quantification

For time-lapse microscopy, embryos were anesthetized in embryo media containing 600μM MS-222 (Sigma) and mounted in 1% low melt agarose (NuSieve GTG). High-resolution time-lapse microscopy of apical protrusions was performed on a Zeiss 880 Airyscan confocal microscope using the fast Airyscan mode and processed using the default parameters. Actin rods were manually counted and quantified in FIJI^29^ using sum slice projections of processed Airyscan image stacks taken at 5 second intervals at the PLLp-skin boundary. All rods quantified were within 4.5μm of the basal surface of the skin (for apical protrusions) or within 4.5μm of the basal surface of the PLLp (for basal protrusions). Bleb fluorescence intensity quantification was performed by measuring the intensity of either LifeAct-Citrine or Lyn-mScarlet along the edge of the bleb in single confocal slices. Intensities were normalized to that of the first image in the blebbing sequence. Retrograde actin flow was quantified by making kymographs along the lamellipodia parallel to the direction of flow.

Segmentation was performed in FIJI by generating regions of interest (ROIs) corresponding to the membrane outline of each cell, followed by manual validation and, if necessary, re-segmentation. These ROIs were converted to 3D models in OBJ format by a macro and models were meshed in Meshlab 2016.12^30^ using the screened poisson surface reconstruction algorithm^31^ followed by simplification by quadratic edge collapse decimation. Rendering was performed in Blender 2.8^32^ using the cycles rendering engine. For long-term timelapse, *Tg(clbnb:lyn-gfp);TgBAC(cxcr4b:h2a-mcherry)* double transgenic embryos at 32hpf were embedded in 1% agarose and imaged on a custom DiSPIM^33^ using a 40X objective after removal of overlying agarose. Image registration and joint deconvolution was performed using a recently improved pipeline that offers greatly increased processing speed^34^. Registration was performed using the methods in Guo *et al*, and for deconvolution we used 10 iterations of conventional Richardson-Lucy. Image rotation was performed using TransformJ^35^ Data analysis was performed in Python 3.7 using the SciPy^36^, Pandas^37^ and NumPy^38^ libraries. Plots were generated in Python using the Matplotlib^39^ and Seaborn^40^ libraries. Raw data in CSV format and Jupyter notebooks containing all statistical analyses and plots are available at https://github.com/chitnislabnih/dallenogare2019

## Results

We set about to image the morphology of individual cells in the PLLp by transplanting cells labelled with a membrane localized eGfp driven by the *cldnb* promoter (Tg(*cldnb:lyn-egfp*)) into *Tg(cldnb:lyn-mscarlet)* transgenic embryos. The resulting chimeric embryos contained mosaically labelled primordia, with a small percentage of donor cells expressing membrane-localized Gfp while the remaining host-derived cells expressed membrane-localized mScarlet (Fig 1A).

**Figure 1.**
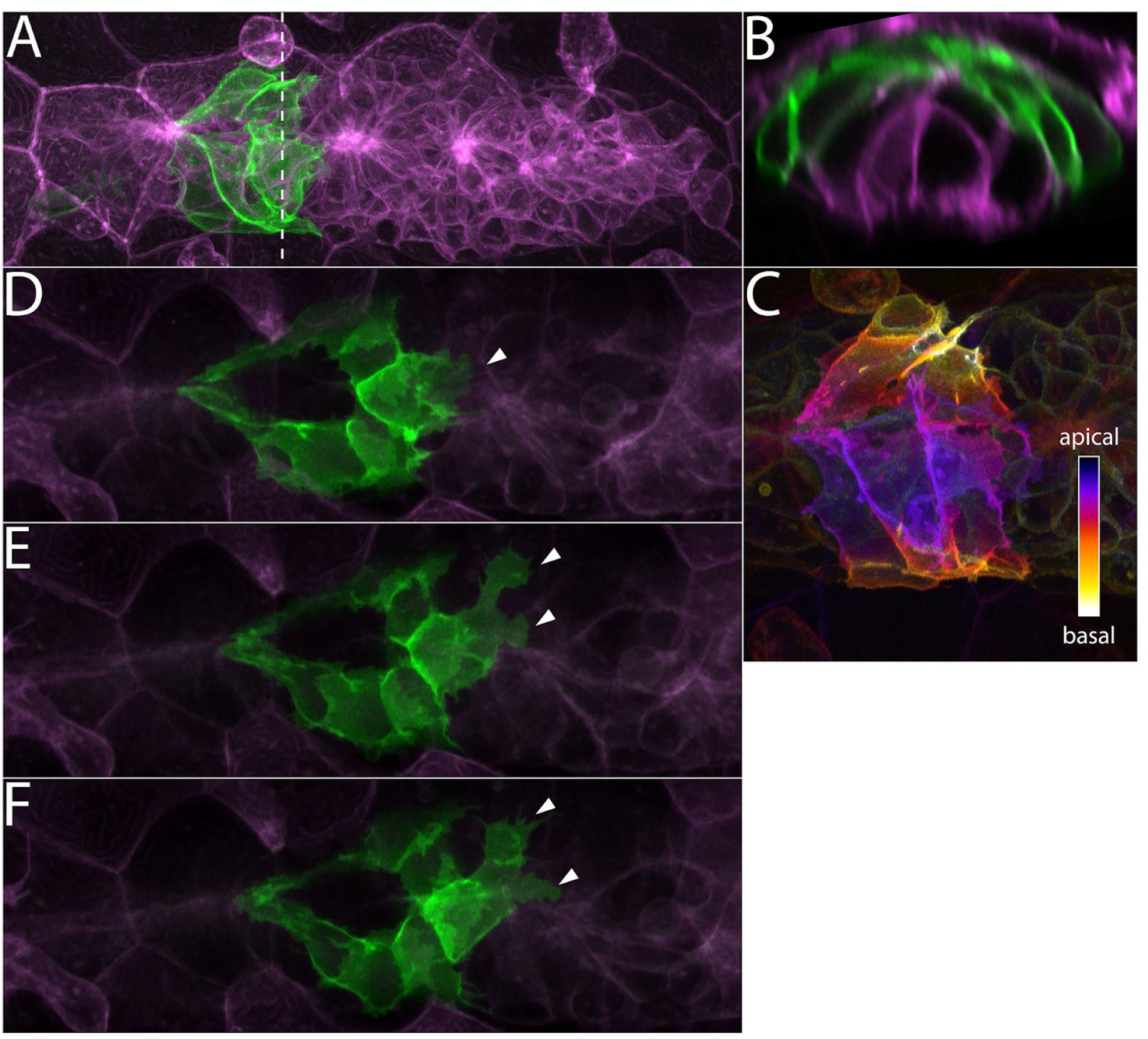
**A**: Z-projection of cells from a *Tg(cldnb:lyn-gfp)* embryo (green) transplanted into a*Tg(cldnb:lyn-mscarlet)* (magenta) embryo. Dashed line indicates the position of the transverse section in B. **B**: Transverse section of A showing position of green transplanted cells. **C**: Apical-Basal depth coding of the *Tg(cldnb:lyn-gfp)* cells shown in A and B. **D-F**: Frames from a timelapse movie showing apical membrane protrusions adjacent to the skin (arrowheads).

In most cases, cells in the trailing domain had the characteristic shape of cells incorporated into an epithelial neuromast -- a basally positioned nucleus and a tightly constricted apical domain that connects with the apical domains of other neuromast cells to form an apical microlumen^41^. In a smaller number of cases, cells in the leading mesenchymal domain were labelled. These cells had a flat morphology without any clear apical-basal polarity, and extended numerous basal membranous protrusions. However, we also observed cells whose cell bodies lie above the level of the neuromast apical constrictions. In some cases, these cells were clearly connected to the apical constriction a nearby neuromast, and they extended laterally across the top of the PLLp and wrapped around the lateral sides (Fig 1B, C). In other cases these apical-dwelling cells had no apparent direct connection to the neuromast apical constriction, often lying between neuromasts. During the course of a timelapse movie, we observed these cells extending multiple broad, flat apical protrusions reminiscent of lamellipodia (Fig 1 D-F, Movie S1). These protrusions were extended closely apposed to the basal surface of the skin, and their similarity to basal cryptic lamellipodia suggested that they might also contribute to the migration of the PLLp.

To further characterize this apical population, we performed high resolution imaging of the PLLp in *Tg(cldnb:lyn-egfp);TgBAC(cxcr4b:h2a-mcherry)* double transgenic fish, in which both the membranes and nuclei of the lateral line primordium are labelled (Fig 2A-D), using DiSPIM microscopy to generate an image with isotropic resolution. While the cells that make up the core of the neuromast have a basally positioned nucleus, the nuclei of these overlying cells resided apically, often above the level of the apical constriction (arrowheads in Fig 2B, C), and directly underneath the skin (Fig 2D). Slicing this image along the coronal plane shows this population of cells occupying a position above the apical constrictions of the neuromast cells (Fig 2B), and sagittal re-sectioning revealed a series of nuclei occupying a circumferential position around the protoneuromast (Fig 2C).

**Figure 2.**
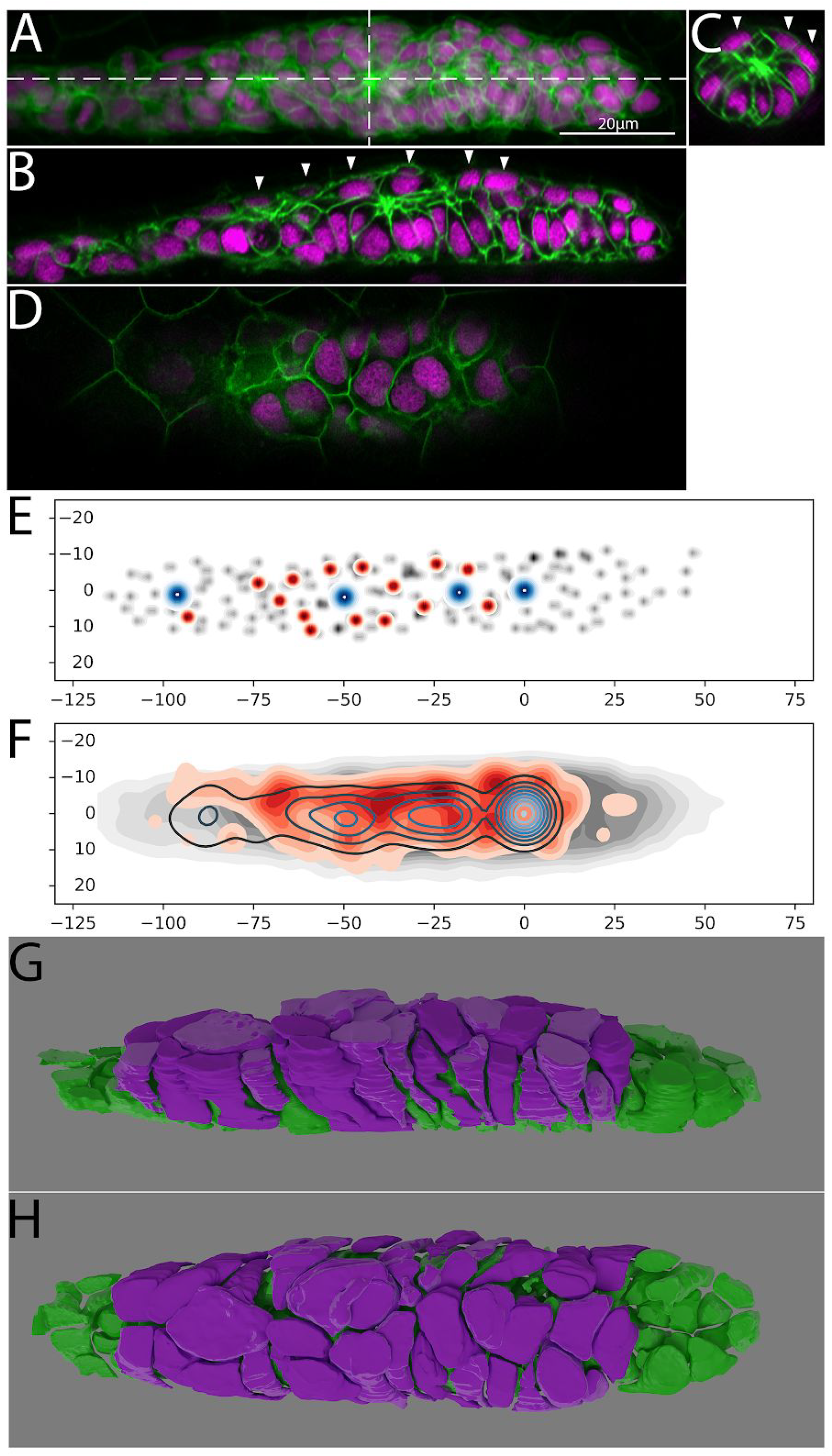
**A**: Z-projection of the PLLp at ~32hpf from individual DiSPIM slices. Dashed lines indicate the position of cross sections in B and C. Membranes are indicated in green and nuclei in magenta. **B**: Sagittal cross section of the PLLp at the position indicated in A. Arrowheads indicate apical nuclei. **C**: Transverse section of the PLLp at the position indicated in A. Arrowheads indicate apical nuclei. **D**: Single confocal slice adjacent to the skin showing apical nuclei. **E**: Schematic of nuclear position in a single PLLp. Grey dots indicate all PLLp nuclei, red dots indicated the position of apical nuclei (see text for details), and blue dots indicate the position of neuromast apical constrictions. **F**: Average of the positions of nuclei in n=10 PLLp. PLLp were aligned based on the position of the leading-most neuromast. Grey indicates the position of all cells, red filled topo-lines indicate the position of apical nuclei and blue open topo-lines indicate the position of neuromast apical constrictions. **G,H**: 3D reconstruction of a confocal scan of a ~32hpf PLLp showing the apical cells in magenta and the remaining cells in green.

To determine the distribution of these cells along the anterior-posterior extent of the PLLp, we mapped the position of all cells whose nuclei resided above the level of the closest neuromast apical constriction and were directly apposed to the overlying skin (red in Fig 2E, F). We then compared their distribution with the distribution of all of the cells of the PLLp (grey in Fig 2E, F) in both single primordia (for example see Fig 2E) and when PLLp were aligned based on the position of the most leading neuromast and the positions aggregated (Fig 2F. These data show that these cells lie above the fully formed neuromasts in the PLLp, but their frequency decreases toward the back of the PLLp, where the trailing-most neuromast is preparing to deposit. However, it should be noted that the apparent lack of these cells in the leading edge is a consequence of our selection criteria, where leading cells, typically flatter and lying closer to the underlying migratory substrate, are excluded. Segmentation and reconstruction of these cells showed that they formed a broad “sheath” that covered the apical side of the PLLp, forming a layer between the apical constrictions of the neuromast and the overlying skin cells. For illustration, Figure 2G, H and movie S2 show a 3D rendering of a fully-segmented PLLp with this population colored in magenta, and the remaining cells are colored in green.

Live imaging to track the position and behavior of these cells over time shows they form a stable population. Apically residing cells could be tracked through more than 3 hours of timelapse during which time they maintained a stable apical position (Fig S1). In these movies we noticed that, consistent with the analysis of the distribution of these cells, they were displaced from their apical positions as the trailing-most neuromast was preparing to deposit, in many cases becoming part of the interneuromast population.

The presence of a cell population with minimal or nonexistent basal contact with the underlying tissue but significant surface area contact with the overlying skin suggested that these cells might contribute to the migration of the PLLp through contacts with the overlying skin. High-resolution imaging suggested the presence of broad membrane protrusions from these cells which were oriented in the direction of migration (Fig 1 D-F). To further examine these protrusions, we performed live imaging using Airyscan super-resolution microscopy in embryos where a BAC containing the *cxcr4b* regulatory elements was used to drive *lifeact-citrine*, which labels F-Actin^42^, in the PLLp. The distribution of LifeAct-Citrine revealed the presence of transient Actin fibers extending within broader membrane protrusions, reminiscent of lamellipodia (Fig 3A, Movie S3). These protrusions were even more apparent when we imaged isolated cells by transplanting cells from *TgBAC(cxcr4b:lifeact-citrine);Tg(cldnb:lyn-mscarlet)* double transgenic embryos into *Tg(cldnb:lyn-mscarlet)* embryos (Fig 3C, movie S4) Quantification of the direction of these protrusions in high resolution timelapse movies of the apical domain of the PLLp showed that they were highly polarized, extending in the direction of migration (Fig 3B), with an average length of ~2.4μm (Fig S2), similar in orientation and length to previously described migratory protrusions that extend from the basal aspect of cells that make up a neuromast. In contrast, the apical surface of the majority of cells in a neuromast are tightly constricted, and we did not observe significant apical protrusive activity from these cells.

**Figure 3.**
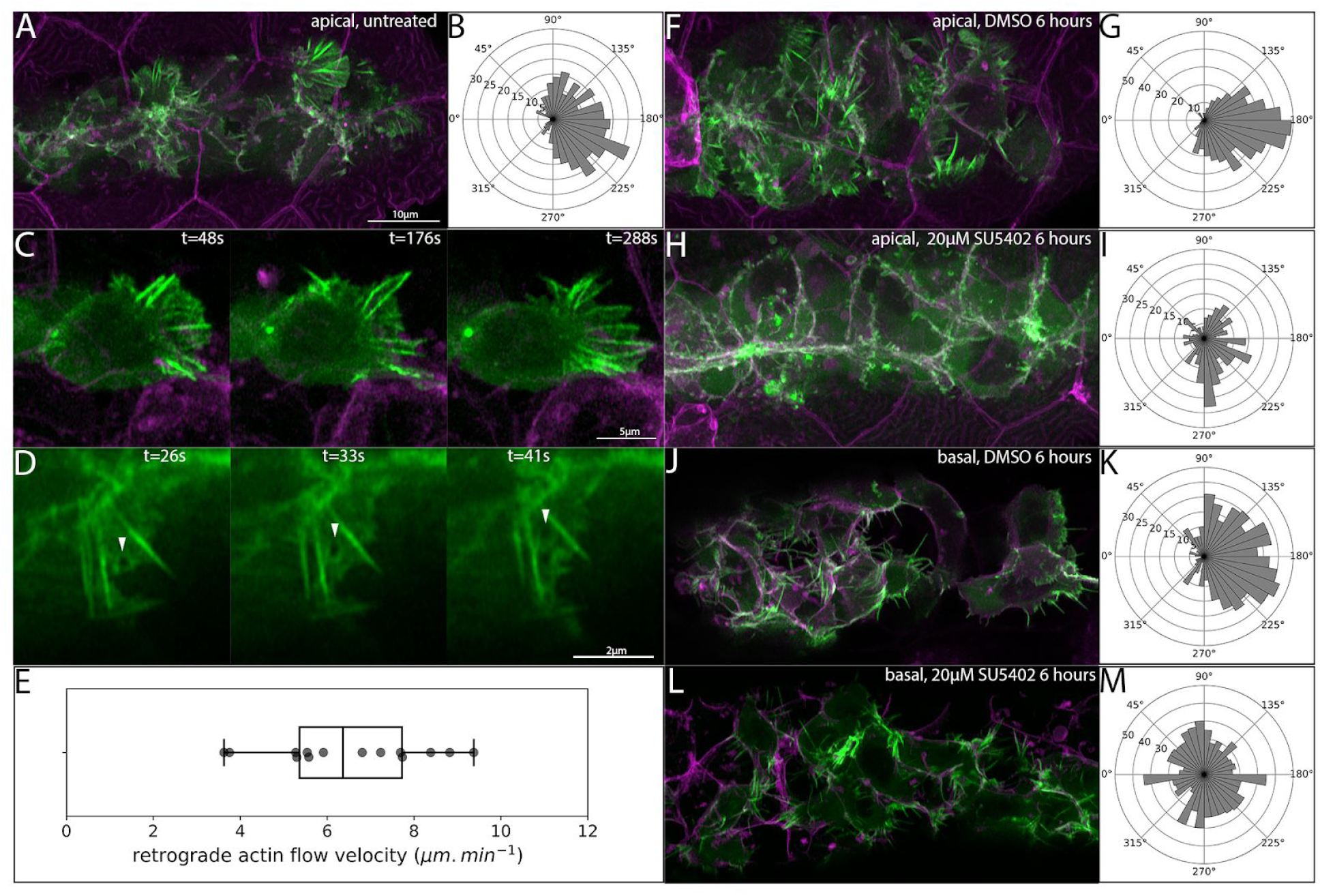
**A**: Maximum intensity projection of a confocal stack showing LifeAct-Citrine-positive projections (green) within 5μm of the skin. **B**: Quantification of the direction of apical actin protrusions from timelapse movies. 180° (right) indicates the normal direction of PLLp migration. **C**: Frames from a timelapse movie showing a single *TgBAC(cxcr4b:lifeact-citrine)* cell adjacent to the overlying skin with polarized protrusive activity. **D**: Series of frames from a timelapse movie showing retrograde flow of LifeAct-Citrine in apical protrusion. **E**: Quantification of retrograde flow velocity in apical protrusions. **F**: Apical LifeAct-Citrine-positive protrusions after 6 hours of treatment with DMSO. **G**: Directionality of apical protrusions after DMSO treatment. **H**: Apical LifeAct-Citrine-positive protrusions after 6 hours of treatment with 20μM SU5402. **I**: Directionality of apical protrusions after SU5402 treatment. **J**: Basal protrusions in *TgBAC(cxcr4b:lifeact-citrine)* transplants after 6 hours of treatment with DMSO. **K**: Directionality of basal protrusions after DMSO treatment. **L**: Basal protrusions in *TgBAC(cxcr4b:lifeact-citrine)* transplants after 6 hours of treatment with 20μM SU5402. **M**: Directionality of basal protrusions after SU5402 treatment. Scale bar for F,H,J,L is the same as A.

In our timelapse movies of *Tg(cxcr4b:lifeact-citrine)* transgenic embryos, we noticed signs of significant retrograde actin flow in these protrusions. In migrating cultured cells, the rate of retrograde actin flow in lamellipodial protrusions has been shown to correlate with the traction force exerted by the migrating cell^43^. To assess the rate of retrograde actin flow, we took high speed super-resolution Airyscan movies at one-second intervals and used the movement of inhomogeneities in the signal to assess the rate of retrograde actin flow in these protrusions (Movie S5). Figure 3D shows one such inhomogeneity flowing backwards from the lamellipodial tip toward the cell body (arrowheads in Fig 3D). Quantification of the flow showed an average retrograde actin flow speed of 6.5μm.min^−1^ with a standard deviation of 1.8μm.min^−1^ (Fig 3E)

The orientation of migratory protrusions on the basal surface of the PLLp is known to be dependent on Fgf signaling, and are lost when Fgf signaling is inhibited by the Fgf receptor inhibitor SU5402^17^. To test whether the orientation of these protrusions, like those on the basal side of the PLLp, is also dependent on Fgf signaling, we treated embryos with either 20μM of SU5402 or DMSO for 6 hours and measured the orientation of the protrusions. After 6 hours of treatment with DMSO, apical protrusions remained robustly polarized in the direction of migration (Fig 3F,G, Movie S6). However, this polarization was lost after Fgf inhibition by SU5402 treatment (Fig 3H,I, Movie S6), suggesting that, like the basal protrusions, the directionality of these apical protrusions was also dependent on Fgf signaling. For comparison, we performed the same analysis on basal protrusions. However, since the densely packed cell membranes comprising the basal surface of the PLLp make quantification of these basal protrusions challenging, we performed these experiments in chimeric embryos where we transplanted *TgBAC(cxcr4b:lifeact-citrine)*;*Tg(cldnb:lyn-mscarlet)* donor cells into *Tg(cldnb:lyn-mscarlet)* host embryos. As expected, in DMSO-treated embryos the basal protrusions were primarily oriented in the direction of migration (Fig 3J, K, Movie S7). However, after 6 hours of treatment with SU5402, this polarity was completely abolished (Fig 3L,M, Movie S7).

The prior experiments demonstrate the existence of a population of cells that make significant contact with the overlying skin during migration. Furthermore, our analysis showed that these cells extended protrusions reminiscent of lamellipodia against the skin, and that the orientation of these protrusions, like that of the basal cryptic lamellipodia, is sensitive to Fgf inhibition. This suggested a potential role for the overlying skin in migration of the PLLp. To test whether the skin is necessary for PLLp migration, we removed the skin overlying the PLLp using a tungsten needle and then imaged the resulting behavior of the PLLp using confocal microscopy. We performed these experiments both in *tg(claudinb:lyn-egfp)* embryos, which labels the periderm layer with membrane localized Gfp, and in *tg(claudinb:lyn-gfp; krt4:dsred)* double transgenic embryos, in which both the periderm and basal layers of the skin are additionally labeled with cytoplasmically localized DsRed^24^. Immediately following skin removal, the PLLp ceased migration and remained stationary.

In most cases, after skin removal, the skin rapidly healed over the PLLp with little to no damage to the PLLp itself (based on the minimal appearance of apoptotic and necrotic cells). In these cases, after skin regrowth, the PLLp recovered normal migration and continued migrating along the length of the embryo (Movie S8). During this subsequent migration neuromast deposition appeared normal, and the primordium reached the tip of the tail after a slight delay (Fig S3). In a minority of cases, where large patches of skin covering a significant fraction of the trunk were removed such that the skin could not heal over the PLLp in the time course of our movies, we observed the PLLp for a period of several hours. In these cases, the PLLp did not recover forward migration, and the PLLp cells eventually underwent apoptosis.

Figure 4A-C shows three still frames from a representative timelapse taken while the skin healed over the PLLp. Initially the PLLp is stationary, having stopped forward migration. The skin (magenta) is healing toward the PLLp (Fig 4A). Twenty minutes later, the skin has made contact with the PLLp, and the morphology of the leading cells has changed, becoming more stretched out, reminiscent of their morphology in intact primordia (Fig 4B). By 97 minutes, the skin has healed over the PLLp and normal forward migration has robustly resumed (Fig 4C). This dramatic behavior can be seen in kymographs of both the green PLLp membranes and the magenta krt4:DsRed-positive epidermal cells (Fig 4D-F). Initially, the PLLp is stationary, while uncovered by the skin (compare figures 4E, F). However, as the skin heals over the PLLp, robust and continuous forward migration is recovered. We removed the skin from >10 independent embryos and, in all cases, migration was abolished until the skin healed over the PLLp.

**Figure 4.**
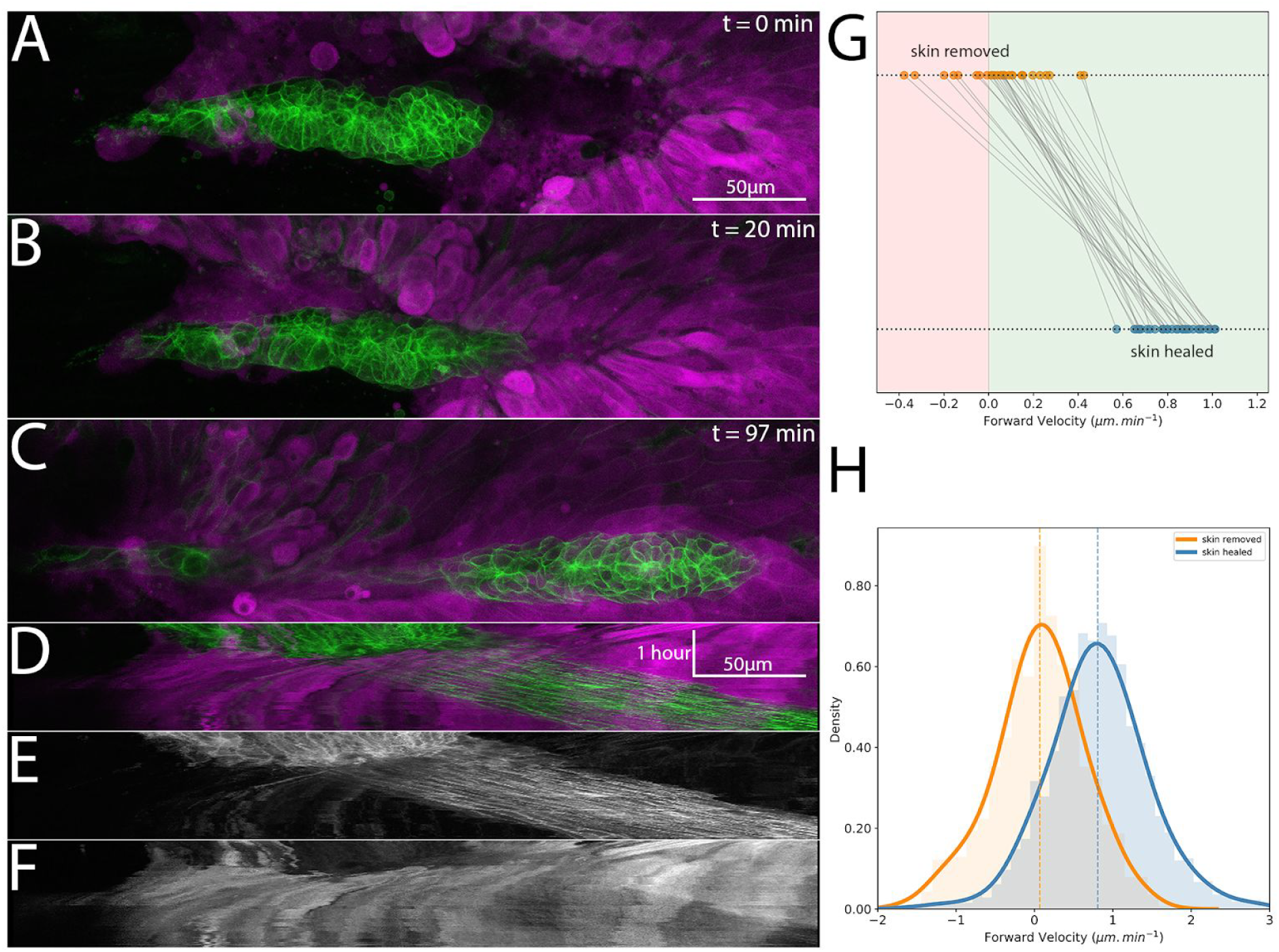
**A-C**: Frames from a timelapse movie showing the skin (basal and periderm layers labelled by *krt4:dsred*) in magenta and the PLLp in green after skin removal and during subsequent healing. **D**: Overlay of kymograph showing movement of the PLLp (green) and healing of the skin over the PLLp (magenta) along the migration course of the PLLp shown in A-C. **E**: Kymograph of the PLLp alone. **F**: Kymograph of the skin alone. **G**: Quantification of the average migration speed of individual cells (n=28) in 3 independent primordia with skin removed (orange dots) and after skin healing (blue dots). Red background indicates rostral movement, green background indicates caudal movement, p=0.0, paired sample t-test. **H**: Histogram of all individual instantaneous cell velocities for the cells measured in G for skin removed (orange) and skin healed (blue). Dashed lines indicate mean velocity for the cells in each condition, and curves indicate the kernel density estimate (kde) for each condition. p=0.0 (unpaired sample t-test)

To quantify this migratory behavior, we crossed *Tg(claudinb:lyn-egfp; krt4:dsred)* fish to *TgBAC(cxcr4b:h2a-mcherry)* fish to visualize the basal cells and periderm of the skin, as well as the both the membranes and nuclei of the PLLp. We then removed the skin overlying the PLLp and tracked the movement of randomly selected cells distributed throughout the PLLp for each of three replicate embryos. For each cell, the time at which the skin covers the position of that individual cell was marked, and the average velocity in the normal direction of migration (along the rostral-caudal axis of the embryo) before and after this point was calculated. Figure 4G shows the paired measurements for each cell, with the top row representing the average velocity of cells before skin contact and the bottom row representing the average velocity after skin contact. Consistent with the results from bulk analysis of movement using kymographs, there was a dramatic increase in forward migration after skin contact. The velocity of cells before skin contact was distributed around 0, suggesting non-directional or “tumbling” movement, and the velocities after skin contact are clustered around a mean of 0.82μm.min^−1^, close to the normal migration speed of a primordium. Aggregating the frame-to-frame velocities shows that the velocity before skin contact is normally distributed around a value of approximately zero (mean = 0.067μm.min^−1^, standard deviation = 0.51μm.min^−1^), wheras the velocities after skin contact are normally distributed around approximately 0.8μm.min^−1^ (mean = 0.807μm.min^−1^, standard deviation = 0.648μm.min^−1^, p=0.00). Taken together, these data suggest a profound inability of the PLLp to migrate without overlying skin, which is completely reversed after skin regrowth over the PLLp.

We hypothesized that removing the skin caused a failure of the overlying cells to extend robust migratory protrusions and that this contributes to the failure of collective migration. To assess this, we crossed *TgBAC(cxcr4b:lifeact-citrine)* fish to *Tg(cldnb4.2:lyn-m-scarlet)* fish to generate double-transgenic embryos in which both Actin and membranes were labelled. The skin over the PLLp was then removed and the resulting Actin dynamics imaged using Airyscan super-resolution confocal microscopy.

When the skin was removed and the PLLp imaged at high spatio-temporal resolution, we noticed a dramatic change in the behavior of the overlying cells. In embryos lacking skin, we saw almost no polarized Actin-rich protrusions of the kind observed extending from overlying cells in intact embryos. Instead, cells appeared disorganized and rapidly extended and retracted protrusions reminiscent of membrane blebs. High-resolution imaging of these bleb-like structures showed that they initiate as a rapid membrane expansion devoid of cortical actin (Fig 5A-D, Movie S9). As the bleb expands, cortical actin is recruited to the newly expanded membrane and can be detected by an increase in the LifeAct-Citrine signal, while the lyn-mScarlet membrane marker remains constant in intensity (Fig 5E). After a short period, these blebs are retracted into the cell.

**Figure 5.**
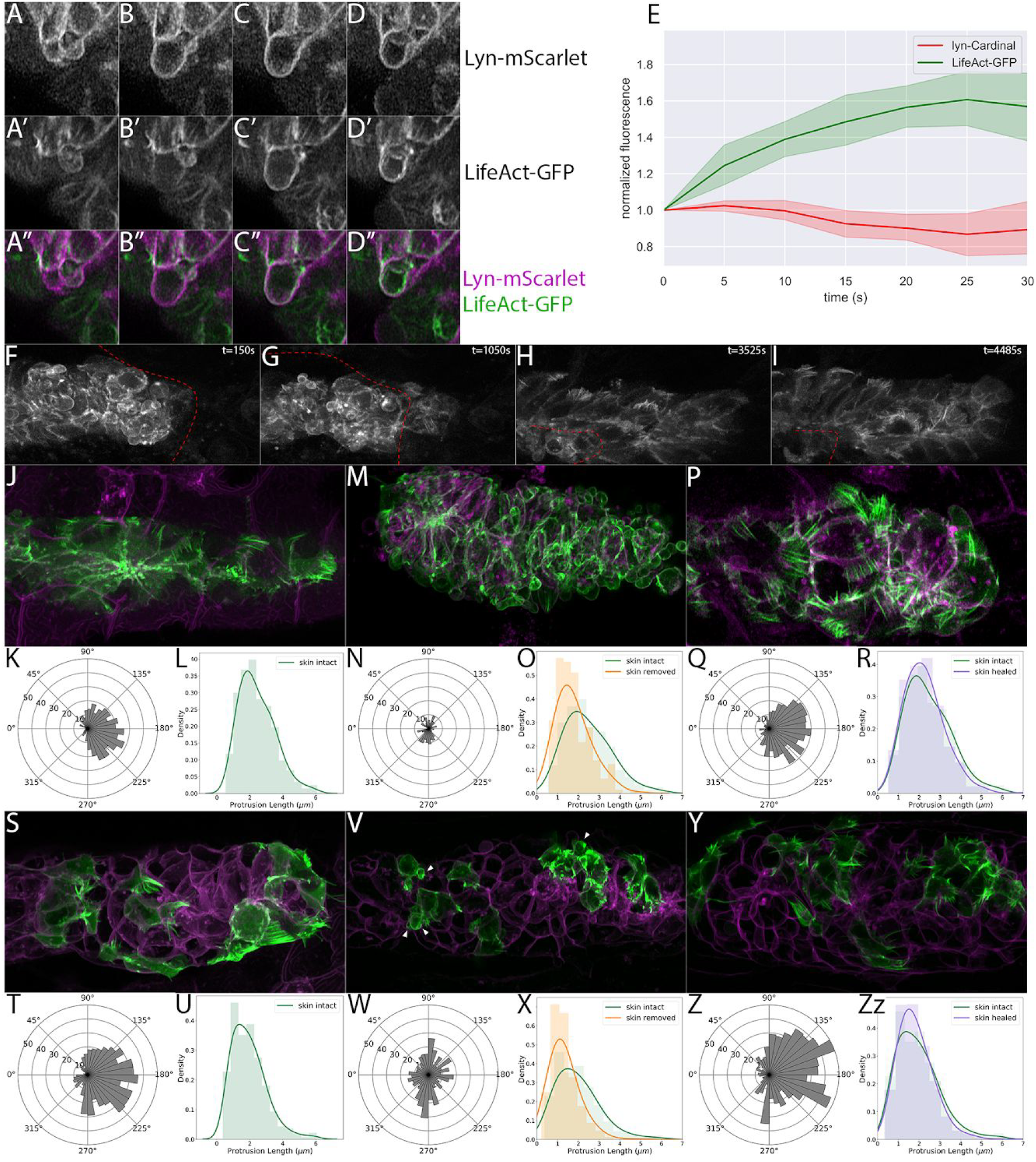
**A-D**: Frames from a timelapse movie showing apical bleb formation and retraction after skin removal. A-D show lyn-mScarlet, marking membranes, A’-D’ show LifeAct-Citrine, marking F-actin and A’’-D’’ show the merge of both channels. **E**: Quantification of membrane (lyn-mScarlet) and Actin (LifeAct-Citrine) along bleb edge over time for n = 8 blebs. Each panel represents an interval of 5s. **F-I**: Frames from a timelapse movie showing apical LifeAct-Citrine during skin healing. Skin position is shown by the dashed red line, and time is indicated in the upper left corner. **J**: Apical LifeAct-Citrine-positive protrusions in embryos with intact skin. **K-L** Quantification of apical protrusion directionality (K) and length (L) in embryos with intact skin. **M**: Apical LifeAct-Citrine after removal of the overlying skin. **N-O**: Quantification of apical protrusion directionality (N) and length (O) in PLLp with overlying skin removed. Apical protrusion length for embryos with intact skin is shown in green in O for comparison. **P**: Apical LifeAct-Citrine after removal and regrowth of the overlying skin. **Q-R**: Quantification of apical protrusion directionality (Q) and length (R) in PLLp with overlying skin removed. Apical protrusion length for embryos with intact skin is shown in green in R for comparison. **S**: Transplanted cells showing basal LifeAct-Citrine-positive protrusions in embryos with intact skin. **T-U**: Quantification of basal protrusion directionality (K) and length (L) in embryos with intact skin. **V**: Transplanted cells showing basal LifeAct-Citrine after removal of the overlying skin. **W-X**: Quantification of basal protrusion directionality (W) and length (X) in PLLp with overlying skin removed. Basal protrusion length for embryos with intact skin is shown in green in X for comparison. **Y**: Basal LifeAct-Citrine after removal and regrowth of the overlying skin. **Z-Zz**: Quantification of basal protrusion directionality (Z) and length (Zz) in PLLp with overlying skin removed. Basal protrusion length for embryos with intact skin is shown in green in Zz for comparison. All images are maximum-intensity projections of confocal stacks within 5μm of the skin (for apical protrusions) or basal surface of the PLLp (for basal slices)

Because the regrowth of the skin over the PLLp is associated with a rapid and robust recovery of forward migration, we imaged Actin dynamics in the PLLp during skin regrowth in *TgBAC(cxcr4b:lifeact-citrine)*; Tg*(cldnb4.2:lyn-m-scarlet)* embryos (Movie S10). After skin removal -- but before skin regrowth over the PLLp -- we observed rapid apical membrane blebbing, as described above. However, as the skin heals over the PLLp, cells undergo a dramatic morphological change. The bleb-type morphology of overlying cells is abolished and cells begin to extend actin-rich protrusions in the direction of migration, reminiscent of those observed apically in intact, unperturbed embryos. This transition was so rapid that over the course of our timelapse movie, we could simultaneously observe blebs from regions of the PLLp not yet covered by skin alongside actin-rich rod-like protrusions from cells that had been covered by the regrowing skin. Eventually the entire PLLp is again covered by the skin, apical protrusive activity is restored, and normal forward migration of the PLLp resumes.

These data suggested a rapid recovery of apical protrusive activity after skin regrowth. To quantify this change, we performed high-resolution imaging of Actin dynamics in *TgBAC(cxcr4b:lifeact-citrine)*;*Tg(cldnb4.2:lyn-m-scarlet)* embryos under three conditions: unperturbed embryos in which the skin has not been removed, embryos in which the skin was removed over the PLLp, and embryos in which the skin had been removed and had subsequently healed over the PLLp. Figure 5 J-R show the results of this experiment. As expected, embryos where the skin had not been removed showed robust protrusions oriented in the direction of migration, with an average length of ~2.4um (Fig 5J-L, Movie S11). When the skin was removed, we again observed a profound loss of apical Actin-rich protrusions, which were replaced by apical membrane blebs of the type shown in figure 5A-D (Movie S12). The few protrusions that remained were both shorter (Fig 5O, compare orange to green, mean intact = 2.45μm, mean skin removed = 1.76μm) and no longer oriented in the direction of migration (Fig 5N). However, as suggested by the timelapse analysis above, after the skin healed over the PLLp, apical protrusions were again observed (Fig 5P, Movie S13), their orientation was again strongly polarized in the direction of migration (Fig 5Q) and their length had recovered to almost unperturbed levels (Fig 5R, compare green and purple, mean intact = 2.45μm, mean post healing = 2.26μm).

Although the absence of these directional apical protrusions was expected after removal of the overlying skin, we were surprised by the profound loss of migratory ability resulting from this manipulation, given that the basal side of the pLLp was presumably still in contact with the underlying tissue. We wondered whether removal of the skin resulted in a broader loss of migratory protrusions in the cells of the PLLp. To assess this, we repeated the above experiment, this time using embryos in which we transplanted donor cells from *Tg*(*BACcxcr4b:lifeact-citrine)* into *Tg(cldnb:lyn-mscarlet)* embryos to generate isolated clones in which we could quantify the directionality and length of basal protrusions.

In intact embryos these basal protrusions, like the apical protrusions, were highly polarized in the direction of migration, although they were on average slightly shorter than apical protrusions (Fig 5 S-U, average length = 1.95μm, Movie S14). When the skin was removed, surprisingly, we observed a profound loss of the normal directional orientation of these basal protrusions (Fig 5W, Movie S15), despite the fact that many of these cells had no contact with the overlying skin. This failure of directional orientation was accompanied, as with the apical protrusions, by a decrease in the average length of the protrusions, (Fig 5X, compare orange to green, mean intact 1.95μm, mean skin removed = 1.32μm) and by the appearance of membrane blebs on the basal surface of these cells (arrowheads in Fig 5V). However, neither the loss of rod-like Actin-rich protrusions nor the appearance of membrane blebs were as dramatic on the basal surface as on the apical surface.

When we imaged these protrusions after skin regrowth (Fig 5Y-Zz, Movie S16), we observed the same dramatic recovery of basal protrusions. The polarization of the basal protrusions in the direction of migration was recovered (Fig 5Z) and their length was similar to that of protrusions in unperturbed embryos (Fig 5Zz, compare purple to green, mean intact = 1.95μm, mean post healing = 1.83μm). Taken together, these data suggest that the profound loss of migratory ability after skin removal does not simply result from loss of apical migratory contacts, but is due to a more general requirement for the presence of an enclosing skin layer in maintaining migration through both apical and basal migratory protrusions.

## Discussion

In this study, we examined the posterior lateral line primordium of Zebrafish, a well-studied model of collective cell migration that migrates in a channel between the underlying somites and overlying epidermis. We show that, in addition to the previously described basal lamellipodia extended by epithelialized cells in this cluster^14^, there is an additional population of cells lying apically, covering these epithelial cells, which also extend migratory processes against the overlying skin. As with the directionality of basal cryptic lamellipodia extended from the epithelial cells that comprise a neuromast, the directionality of these basal processes is abolished by inhibiting Fgf signaling by treatment with SU5402, suggesting that the requirement for Fgf signaling to maintain migration applies to both apical and basal processes. Although this study has focused on the apical protrusions extended by these cells against the overlying skin, we also note that this population - while relatively stable - can also displace and move to the lateral edges of the primordium. This suggests the possibility that migratory activity is not confined to the basal or apical surfaces of the primordium but in a circumferential sheath that surrounds the central core neuromasts.

Typically, the PLLp has been conceptually separated into a leading mesenchymal-like domain, where cells are relatively flat and have no obvious apical-basal polarity, and a trailing epithelial domain, where cells become elongated and adopt a distinct epithelial morphology with apical-basal polarity. Despite this distinction, it has long been recognized that both domains contribute to collective migration, with epithelial cells extending cryptic lamellipodia in the direction of migration. In this study, we have defined the “sheath” cells as those that occupy an apical position above the neuromasts, by definition excluding leading mesenchymal cells. However, it is possible that, rather than being an entirely separable population, this “sheath” represents and extension of non-epithelial cells over the entire PLLp and these cells constitute a continuous migratory population. In this context, rather than conceptually dividing the PLLp into a leading and trailing domain with distinct cellular morphologies, a second axis is radially arranged with apico-basally polarized epithelial cells in the core and more mesenchymal cells at the periphery, an example of what Blanchard and co-authors have called “mesoscale heterogeneity” in migrating systems^44^.

Consistent with the existence of migratory contacts on the apical side of the PLLp closely apposed to the overlying skin, we show that when the skin is removed apical lamellipodia are almost completely abolished and cells extend and retract rapid membrane blebs. The few Actin-rich rod-like protrusions that remain are no longer oriented in the direction of migration and are significantly shorter than those extended from the apical surface of the PLLp when the skin is intact. Regrowth of the skin restores both the normal length and directionality of these protrusions with very little delay suggesting, again, a rapid switch in migratory ability and morphology between confined and unconfined PLLp cells.

Intriguingly, we observe very similar changes in the basal cryptic lamellipodia of neuromast cells when the skin is removed, despite the fact that these cells, in many cases, do not directly contact the overlying skin. While Actin-rich protrusions are still observed at the basal surface of the PLLp these protrusions, like those observed apically, become shorter and are no longer oriented in the direction of migration. Concurrent with this loss of directionality, we see the appearance of membrane blebs in the basal surface of these cells, although to a lesser extent than is observed apically. This observation, along with the rapid recovery of directionality and length of basal protrusions after skin regrowth--despite the fact that these cells to not themselves directly contact the skin--raises the possibility that the skin is not only necessary to provide a substrate for the apical migratory processes, but that it is necessary to provide some confinement in order for productive migratory processes to be extended from both the basal and apical surfaces.

This rapid switch between distinct cell morphologies has also been reported in early zebrafish cells placed in culture^45^. When in culture, these cells show a characteristic but unproductive blebbing, and fail to migrate. However, upon either treatment with Lysophosphatidic Acid or by being subject to mechanical confinement, the cells adopted a pear-shaped morphology with a large stable bleb and became highly migratory. Intriguingly, when the confining substrate is non-adhesive, the migratory ability of these cells increases, suggesting that this migratory mode does not rely on specific adhesions with the confining substrate. This migration was associated with rapid retrograde cortical actin flow in the leading edge of the cell. Whether or not these superficial similarities reflect deep mechanistic similarities in the mechanisms of migration in these two contexts will require further study.

In other contexts, changes in adhesion and confinement can lead to rapid changes in cell morphology in ways that are less consistent with what we observe in the PLLp. Bergert *et al*^46^ placed Walker 256 carcinosarcoma cells in confinement and showed that a rapid switch from a primarily bleb-based migration mode and a mixed bleb-lamellipodia mode can also be induced by increasing substrate adhesion. This switch could be rapidly reversed when cells moved onto a non adhesive surface, with cells no longer forming lamellipodia and reverting to a primarily bleb-based migratory mode. The interplay between adhesion, confinement, and cortical contractility has been explored systematically by Liu and colleagues^47^, who suggest that these three parameters integrate to determine the mechanism of migration. Similarly, in Dictyostelium, confinement itself is sufficient to induce rapid changes between blebbing and pseudopod-mediated migration^48^. In this context, reducing confinement by increasing the height of a microfluidic channel through which the cells are migrating is associated with a rapid loss of blebs and an increase in pseudopod formation. However it should be noted that in these other contexts cells can switch to bleb formation as a productive mode of migration, whereas in the PLLp this transition is associated with non productive blebs and a failure of migration.

How which cells migrate through a confined environment is poorly understood, especially during collective migration. Unconfined cells in 2D culture can migrate by promoting attachment to a substrate through which force can be transmitted. However, in confined systems such as 3D scaffolds, the mechanical properties of the environment itself can force cells into significant contact with their extracellular migratory substrate, allowing the transmission of traction forces without adhesion (for review, see Paluch et al, 2016^49^). This non-adherent migration has been demonstrated in a variety of contexts^47,50^ and has been proposed to allow for efficient movement of cells through complex environments without the energetic cost of continuously forming and breaking attachments. In the context of the migrating PLLp it is likely that the overlying protrusions are directly contacting the cells of the overlying skin, where classical integrin-based adhesions might not be desirable or possible. The fact that in the vast majority of cases (>95%), we were able to remove the skin from the PLLp while retaining attachment of the PLLp to the underlying substrate suggests an asymmetry of adhesion between the apical and basal surfaces of the PLLp.

So-called “chimneying” mechanisms, whereby cells can push outwards against the surrounding environment and generate enough force to allow migration, have been suggested to play a role in migration of leukocytes in an adhesion-independent manner^51^. A related but mechanistically distinct mechanism, flow-friction-driven force transmission, is hypothesized to transmit intracellular force from the cytoskeleton to the substrate by means of nonspecific friction between the cell and its environment^45,52^. In this context retrograde flows of the actomoysin cortex, similar to those we observe in apical protrusions in the PLLp, have been suggested to play a role in the generation of motile force^49^. Gardel and colleagues^43^ have demonstrated a biphasic relationship between retrograde flow speed and traction, with traction and flow directly correlated at speeds of below 10nm.s^−1^ (~0.6μm.min^−1^) but inversely correlated at higher speeds. The retrograde flow speeds we measure are relatively fast, on the order of 6.5μm.min^−1^ approximately an order of magnitude faster than the limit for enhancing traction stress measured by Gardel *et al* and ~6 fold higher than the average migration speed of the PLLp itself. The relatively rapid retrograde actin speed in lamellipodia of the lateral line is consistent with a low-adhesion migratory mode, as adhesions are thought to couple the intracellular actin mesh to the ECM and slow the retrograde flow of actin via a molecular “clutch” mechanism^53^. However, future experiments will be necessary to examine the relationship between adhesion and retrograde actin flow in this system.

Understanding how cells in the developing embryo move through diverse environments is critical for understanding both normal morphogenesis and a number of pathologies. Cell migration has been studied extensively in cell culture in the two-dimensional regime, however there is growing evidence that 2D models emphasize specific modes and mechanisms of cell migration that might not apply to cells migrating in a three-dimensional confined environment. While multicellular systems acquire behaviors that correspond broadly to those observed in single cells, it is likely that the details about how they acquire those characteristics differ significantly. The development of in vivo model systems to study cell migration at high spatio-temporal resolution, as well as quantitative methods for analyzing such systems will be necessary to extend the study of 3D cell migration from the dish to the animal.

## Acknowledgements

The authors wish to thank Holger Knaut and Alvaro Sagasti for generously sharing transgenic lines and Angela Hvitved for editing the manuscript. Portions of this work were performed in collaboration with the Advanced Imaging and Microscopy (AIM) Resource at the NIBIB. This work was supported by the intramural research program of the Eunice Kennedy Shriver National Institute of Child Health and Human Development at the National Institutes of Health (HD001012). All work was performed in accordance with animal study protocol 18-013

---

**Figure S1.**
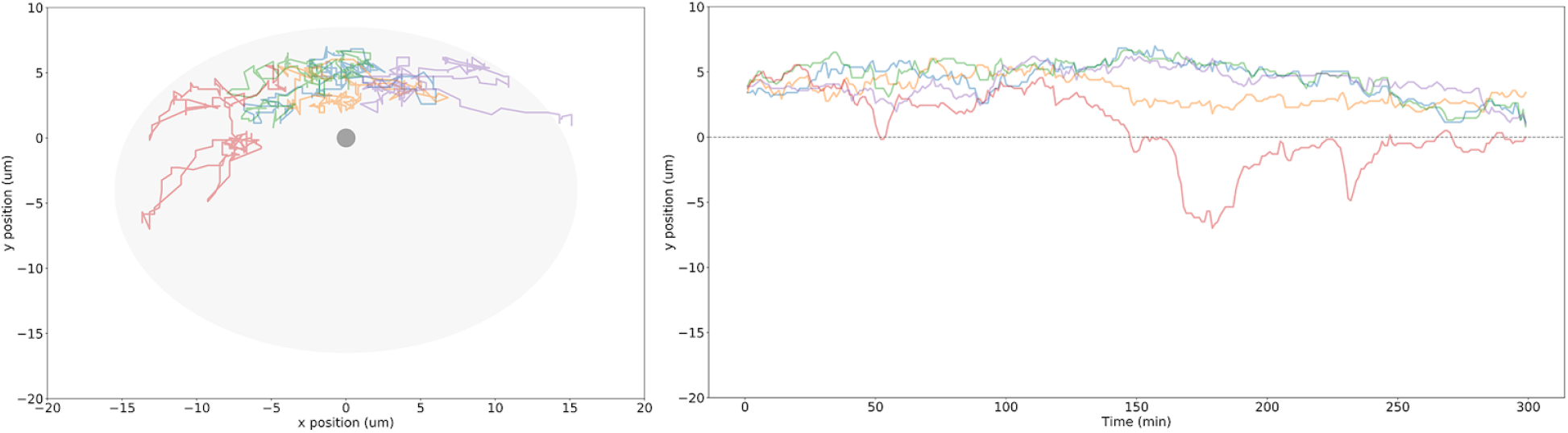
Tracks for 5 cell nuclei over 300 minutes of timelapse. Left panel shows cross section view of tracks, with the central grey dot at (0,0) indicating the position of the closest neuromast, and the grey outline the approximate shape of the PLLp. Right panel shows side view of tracks. y-axis represents the apical-basal position of the cell nucleus (with 0 indicating the level of the neuromast apical constriction), and the x-axis represents time. Track colors are the same in both plots.

**Figure S2:**
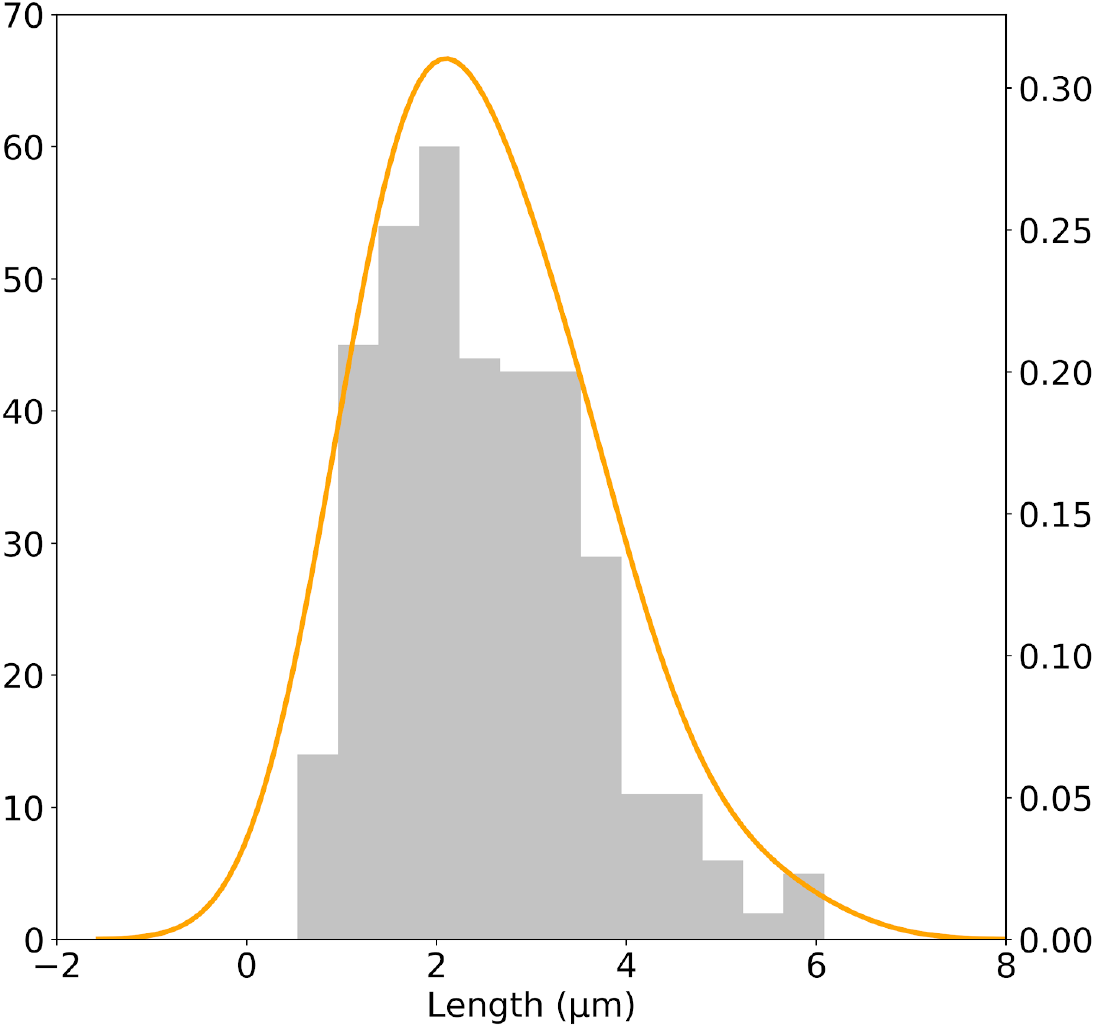
Distribution of lengths of apical actin filaments

**Figure S3.**
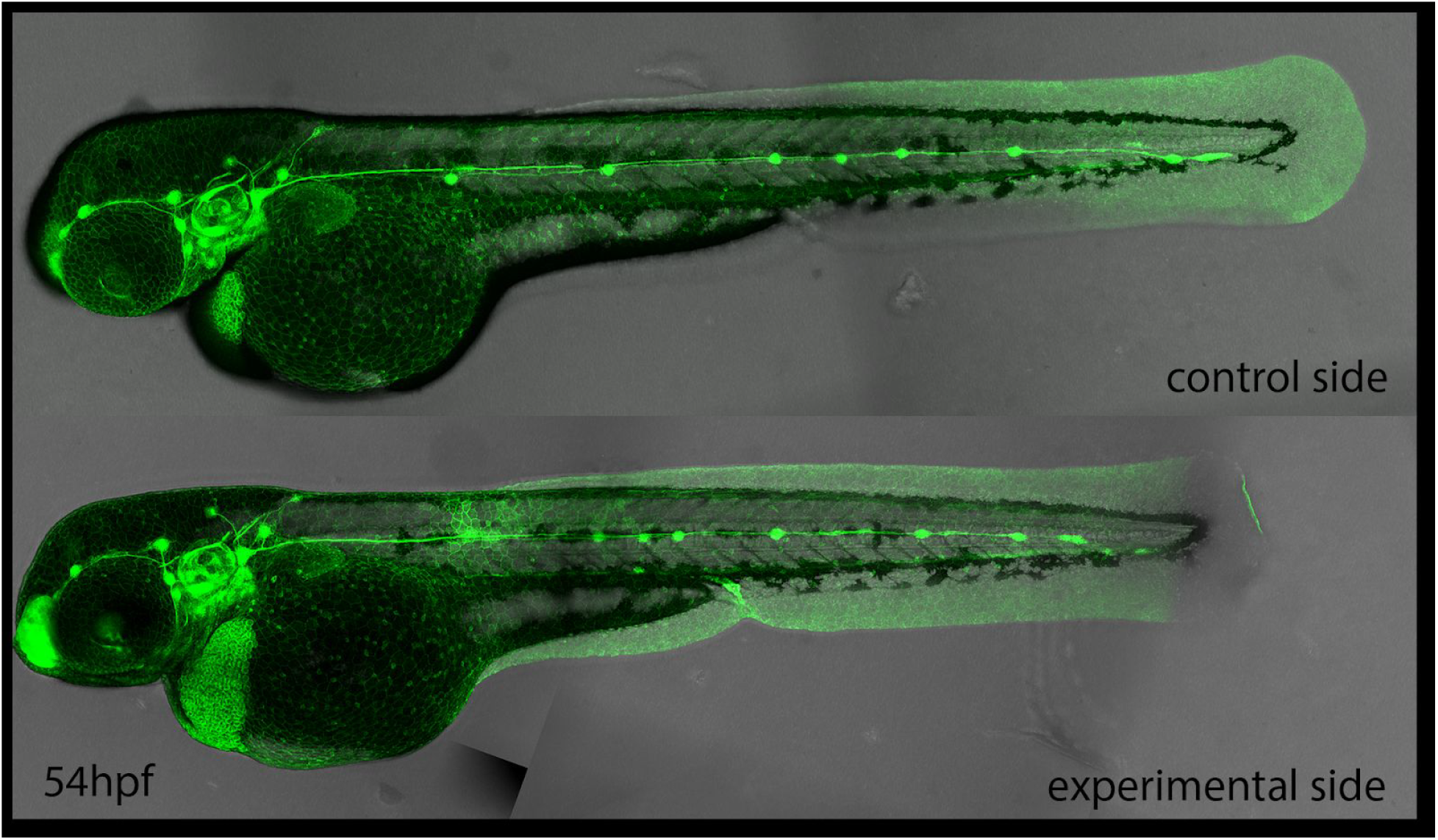
Image of an embryo where skin had been removed over the PLLp at ~32hpf and migration had been allowed to proceed until 54hpf. Top panel shows control (unmanipulated) and bottom panel shows experimental (skin removed) side of the same embryo.

Movie S1. Related to figure 1. Z-projection of a timelapse movie of *Tg(cldnb:lyn-gfp) (green)* cells transplanted into *Tg(cldnb:lyn-mscarlet)* embryos, showing apical membrane protrusions.

Movie S2. Related to figure 2. 3D rendering of a reconstruction of the PLLp from confocal data showing apical cells colored in magenta and remianing PLLp cells colored in green.

Movie S3. Related to figure 3. Z-projection of a timelapse movie taken adjacent to the skin in *TgBAC(cxcr4b:lifeact-citrine)* (green); *Tg(cldnb:lyn-mscarlet)* (magenta) double transgenic embryos, showing apical protrusive activity.

Movie S4. Related to figure 3. Z-projection of a timelapse movie taken adjacent to the skin in an embryo where *TgBAC(cxcr4b:lifeact-citrine)* (green); *Tg(cldnb:lyn-mscarlet)* (magenta) double transgenic cells have been transplanted into *Tg(cldnb:lyn-mscarlet)* embryos, showing apical protrusive activity from a single cell.

Movie S5. Related to figure 3. Z-projection of a timelapse movie taken adjacent to the skin in *TgBAC(cxcr4b:lifeact-citrine)* embryos, showing rapid retrograde actin flow in the apical protrusion.

Movie S6. Related to figure 3. Z-projection of a timelapse movie taken adjacent to the skin in a *TgBAC(cxcr4b:lifeact-citrine)* (green); *Tg(cldnb:lyn-mscarlet)* (magenta) double transgenic embryo. Top panel shows an embryo which has been treated with DMSO for 6 hours, bottom panel shows an embryo which has been treated with 20μM SU5402 for 6 hours.

Movie S7. Related to figure 3. Z-projection of a timelapse movie taken of basal protrusions in a *TgBAC(cxcr4b:lifeact-citrine)* (green); *Tg(cldnb:lyn-mscarlet)* (magenta) double transgenic embryo. Top panel shows an embryo which has been treated with DMSO for 6 hours, bottom panel shows an embryo which has been treated with 20μM SU5402 for 6 hours.

Movie S8. Related to figure 4. Z-projection of a timelapse movie of *Tg(cldnb:lyn-gfp)* (green)*; Tg(krt4:dsred)* (magenta) double transgenic embryos in which the skin had been removed over the PLLp and allowed to heal, showing failure of migration when uncovered by skin and subsequent recovery of migration after skin healing.

Movie S9. Related to figure 5. Z-projection of a timelapse movie of *TgBAC(cxcr4b:lifeact-citrine)* (green); *Tg(cldnb:lyn-mscarlet)* (magenta) embryos showing blebbing of apical cells after skin removal.

Movie S10. Related to figure 5. Z-projection of a timelapse movie of *TgBAC(cxcr4b:lifeact-citrine); Tg(krt4:dsred)* (omitted for clarity) double transgenic embryos showing the disappearance of apical blebs and the reappearance of apical lamellipodia-like protrusions after skin healing. Red line indicates the position of the healing skin marked by the position of the *Tg(krt4:dsred)* signal.

Movie S11. Related to figure 5. Z-projection of a timelapse movie taken adjacent to the skin in an intact *TgBAC(cxcr4b:lifeact-citrine)* (green); *Tg(cldnb:lyn-mscarlet)* (magenta) double transgenic embryo, showing apical protrusive activity.

Movie S12. Related to figure 5. Z-projection of a timelapse movie taken adjacent to the skin in a *TgBAC(cxcr4b:lifeact-citrine)* (green); *Tg(cldnb:lyn-mscarlet)* (magenta) double transgenic embryo where the skin had been removed over the PLLp, showing apical blebbing.

Movie S13. Related to figure 5. Z-projection of a timelapse movie taken adjacent to the skin in a *TgBAC(cxcr4b:lifeact-citrine)* (green); *Tg(cldnb:lyn-mscarlet)* (magenta) double transgenic embryo where the skin had been removed over the PLLp and allowed to subsequently heal over the PLLp.

Movie S14. Related to figure 5. Z-projection of a timelapse movie taken of basal protrusions in an intact embryo where *TgBAC(cxcr4b:lifeact-citrine)* (green); *Tg(cldnb:lyn-mscarlet)* (magenta) double transgenic cells had been transplanted into *Tg(cldnb:lyn-mscarlet)* (magenta) single-transgenic embryos.

Movie S15. Related to figure 5. Z-projection of a timelapse movie taken of basal protrusions and blebs in an embryo where *TgBAC(cxcr4b:lifeact-citrine)* (green); *Tg(cldnb:lyn-mscarlet)* (magenta) double transgenic cells had been transplanted into *Tg(cldnb:lyn-mscarlet)* (magenta) single-transgenic embryos and the skin removed over the PLLp.

Movie S16. Related to figure 5. Z-projection of a timelapse movie taken of basal protrusions and blebs in an embryo where *TgBAC(cxcr4b:lifeact-citrine)* (green); *Tg(cldnb:lyn-mscarlet)* (magenta) double transgenic cells had been transplanted into *Tg(cldnb:lyn-mscarlet)* (magenta) single-transgenic embryos, the skin removed over the PLLp and allowed to subsequently heal over the PLLp.

